# Museum specimens of a landlocked pinniped reveal recent loss of genetic diversity and unexpected population connections

**DOI:** 10.1101/2022.05.19.492422

**Authors:** Matti T. Heino, Tommi Nyman, Jukka U. Palo, Jenni Harmoinen, Mia Valtonen, Małgorzata Pilot, Sanni Översti, Elina Salmela, Mervi Kunnasranta, A. Rus Hoelzel, Minna Ruokonen, Jouni Aspi

## Abstract

**Aim:** The Saimaa ringed seal (*Pusa hispida saimensis*) is endemic to Lake Saimaa in Finland. The subspecies is thought to have originated when parts of the ringed seal population of the Baltic region were trapped in lakes emerging due to post-glacial bedrock rebound around 9,000 years ago. During the 20^th^ century, the population experienced a drastic human-induced bottleneck. Today encompassing a little over 400 seals with extremely low genetic diversity, it is classified as endangered. Our main aim was to evaluate the role of the 20^th^ century bottleneck in the erosion of genetic diversity in the Saimaa seal population. We also evaluated connections with other ringed seals from the Baltic Sea, Lake Ladoga, North America, Svalbard and the White Sea.

**Location:** Lake Saimaa, Finland, together with the Baltic Sea and the Arctic Ocean.

**Methods:** We sequenced sections of the mitochondrial control region from 60 up to 125 years old museum specimens of the Saimaa ringed seal. The generated dataset was combined with publicly available sequences. We studied how genetic variation has changed through time in this subspecies, and how it is phylogenetically related to other ringed seal populations.

**Results:** We observed temporal fluctuations in haplotype frequencies and loss of haplotypes accompanied by a recent reduction in female effective population size. In apparent contrast with the traditionally held view of the Baltic origin of the population, the Saimaa ringed seal mtDNA variation shows also affinities to North American ringed seals.

**Main conclusions:** Our results suggest that the Saimaa ringed seal has experienced recent genetic drift associated with small population size. The results further suggest that extant Baltic ringed seals do not represent well the ancestral population of the Saimaa ringed seal, which calls for re-evaluation of the deep history of this subspecies.

## 1 INTRODUCTION

Modern DNA sequencing technologies and analytical methods, especially simulation-based approaches, enable detailed molecular inferences on population-level processes and demographic history in human and wildlife populations (Foote et al., 2021; Hoban et al., 2012; Taylor et al., 2021; Vilaça et al., 2021). However, there are limits as to what can be deduced from present-day genetic data. This is largely since the contemporary genetic composition of any population has been shaped by the actions of all evolutionary forces (drift, mutation, migration and selection) throughout the whole history of the population. Understanding the population dynamics at a certain period would require disentangling the effects of these forces, which is often impossible. The confounding signals arising from the actions of the different forces at different times can be to some extent minimized, for example, by marker choice (ignoring selection, assuming steady mutation rates), but drift and gene flow are harder to partition.

These limitations of contemporary variation can nevertheless be circumvented by directly obtaining data on past diversity. Temporal sampling of varied genetic data, *e*.*g*., from subfossil, archaeological or museum specimens, can reveal the developments leading to the palimpsest of current genetic diversity and provide a substantially more accurate view of past evolutionary history. In this context, zoological museum collections have proven an invaluable source of direct knowledge on temporal changes in DNA diversity and, ultimately, on the demography of various wildlife species (Hofreiter et al., 2015; Huynen et al., 2012; Nakahama, 2021; Rizzi et al., 2012).

The Saimaa ringed seal (*Pusa hispida saimensis*) offers an excellent model population for exploring the effects of small size and isolation on genetic diversity. Endemic to Lake Saimaa in Finland (Fig. 1), its origin and timing of isolation are deemed as well established. It is considered a subspecies of the Holarctic ringed seal, which, numbering in millions is the most abundant Arctic pinniped (Reeves, 1998). Based on the understanding of the geological history of the Baltic Sea region, the population originates from marine ringed seals that colonized the Baltic basin during the deglaciation of the Scandinavian Ice Sheet, c. 10,000 years ago (Ukkonen, 2002; Ukkonen et al., 2014). In consequence of post-glacial bedrock rebound, parts of the Baltic ringed seal population (*currently P. h. botnica*) were trapped in emerging freshwater lakes, including lakes Saimaa and Ladoga (subspecies *P. h. ladogensis*). Hence, the Saimaa ringed seal has lived in complete isolation for c. 860 generations (Palo et al., 2003), during which it has evolved into a morphologically, behaviorally and genetically distinct subspecies (Berta & Churchill, 2012; Hyvärinen & Nieminen, 1990; Kunnasranta, 2001). Today, the Saimaa ringed seal population encompasses c. 420–430 seals (Metsähallitus, 2020) and is classified as endangered, both nationally (Liukko et al., 2016) and internationally (Kunnasranta et al., 2021). During the 20^th^ century, the population experienced a drastic human-induced bottleneck from hunting and later due to environmental pollutants.

**Figure 1.**
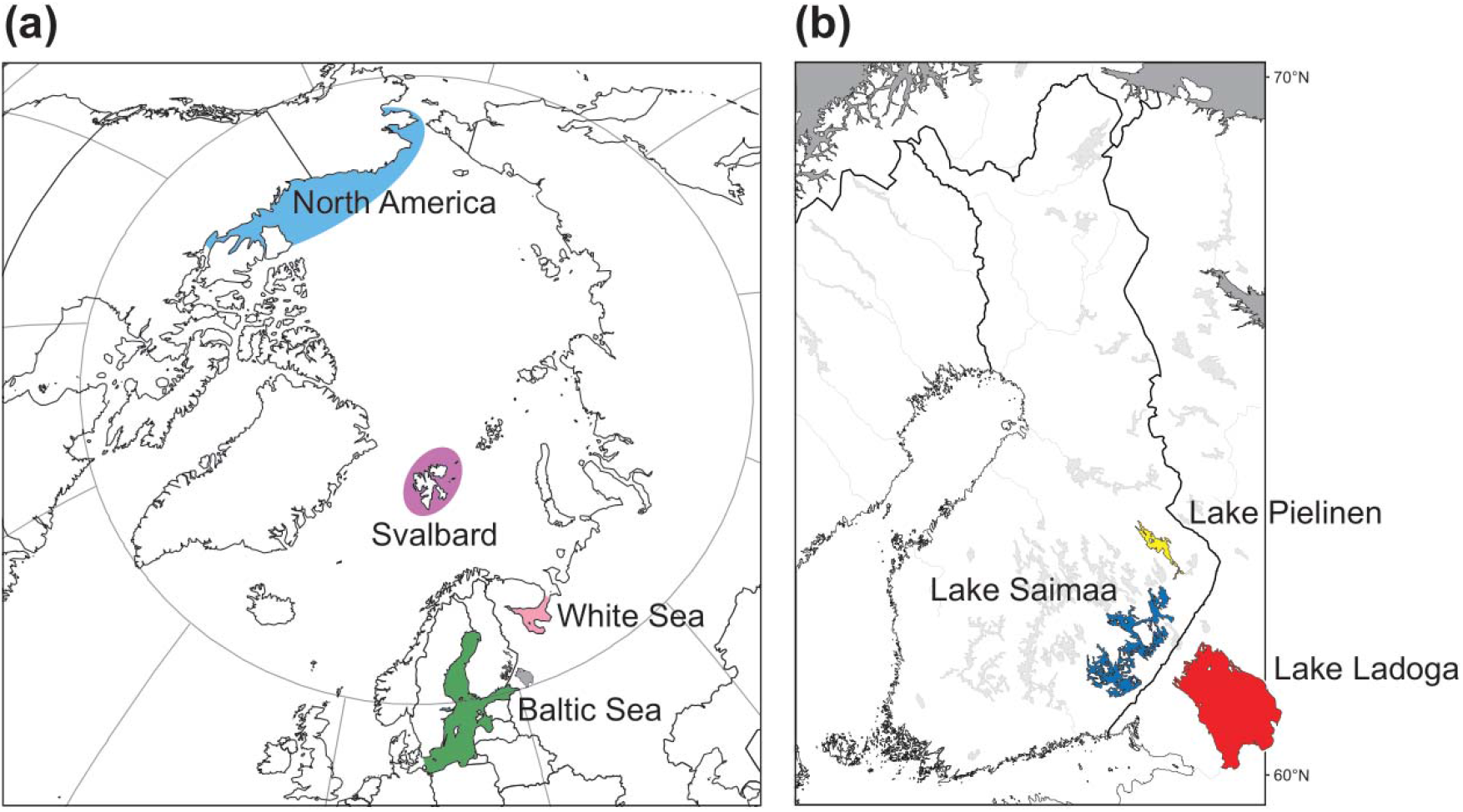
Maps showing the areas from which (a) marine, and (a) freshwater ringed seals were sampled for this study. (Note that a permanent population is not present in Lake Pielinen).

Due to its limited lacustrine habitat, the population has probably never been large; based on the size and resources of the lake, the maximum number of seals has been estimated as being few thousands (Hyvärinen et al., 1999). In the late 1800s, there were possibly as many as 1,000 seals remaining in Lake Saimaa, as suggested by bounty statistics (Kokko et al., 1999) and local knowledge (Hyvärinen et al., 2004). The dwindling of the population started in the early 1900s and continued through most of the century despite protection of the subspecies in 1955. In the 1980s, the population reached its ultimate low of fewer than 150 seals (Sipilä et al., 1990). Since then, intensified conservation efforts (aiming at *N* > 400) set the population to a slow increase in the 1990s, and the positive trend has now continued for more than 20 years (Kunnasranta et al., 2021).

In comparison with other ringed seal subspecies – and almost any mammal population –, genetic diversity of the Saimaa ringed seal is extremely low (Martinez-Bakker et al., 2013; Palo et al., 2003; Valtonen et al., 2012, 2014). Roughly 55% of the autosomal heterozygosity and 90% of the mitochondrial nucleotide diversity present today in the marine population has been lost during the founder event and isolation. The diversity is also substantially lower than in Ladoga ringed seals (Nyman et al., 2014; Valtonen et al., 2012, 2014). Although the Saimaa ringed seal is known to have suffered a substantial reduction in numbers during the 20 ^th^ century, it is not known how much of the total reduction of genetic diversity can be attributed to these recent events. Previous genetic studies have estimated a population trajectory spanning back to the isolation timepoint (Nyman et al., 2014; Palo et al., 2003; Valtonen et al., 2012), but have suggested a minor role for the 20^th^ century bottleneck in the total diversity loss. Also, Nyman et al. (2014), exploring the plausible demographic histories leading to the current diversity using Approximate Bayesian Computation (ABC) analyses, could not find conclusive support for models including the recent bottleneck, over models including only an ancient bottleneck associated with the colonization of Lake Saimaa.

Using mitochondrial DNA (mtDNA) control-region sequences obtained from 60 up to 125 years old museum specimens, as well as previously published contemporary data, we explored changes in Saimaa ringed seal mtDNA diversity. Our main aim was to evaluate whether the loss of genetic variation of the Saimaa ringed seal can be primarily attributed to slow erosion during its isolation after a founder event, as previously suggested (Nyman et al., 2014; Palo et al., 2003; Valtonen et al., 2012), or to the 20^th^-century bottleneck (Peart et al., 2020; Stoffel et al., 2018). We also evaluate the history of the Saimaa population by positioning the diversity in Saimaa on the cladistics canvas of mtDNA variation found in ringed seals from the Baltic Sea, Lake Ladoga, North America (Chukchi and Beaufort Sea), Svalbard and the White Sea.

## 2 METHODS

### 2.1 DNA extraction, amplification and sequencing

We obtained tissue samples (bones, claws or skin) of 63 historical Saimaa ringed seals from the collections of the Finnish Museum of Natural History (Supplementary Table S1). DNA extractions, amplifications and sequencing were performed either at the University of Oulu, Finland, or at Durham University, U.K.

At the University of Oulu, DNA was extracted in the ancient-DNA laboratory of the Centre for Material Analysis, following the protocol by Rohland et al. (2010). For bones and claws, a small portion of the surface of each sample was removed, and circa 250 mg of powder was obtained by drilling into the sample; the powder was then subjected to DNA extraction. For hides, a circa 250 mg piece was cut into small pieces with a sterile scalpel and then subjected to extraction. We targeted a circa 704-bp section of the mitochondrial control region (with some length variation observed among the samples). The whole region was amplified and sequenced in short variable-sized overlapping fragments depending on sample quality (for primers, see Supplementary Table S2). We first attempted to amplify each sample using primer pairs H16305/R328, F322/R471 and F425/L224. If this failed, amplification of shorter fragments was performed using primer pairs H16305/R109, F108/R328, F322/R471, F425/R605 and F603/L224. Each PCR included 1X AmpliTaq Gold® 360 Buffer, 2 mM of MgCl2, 1 mg/ml of BSA, 200 µM of each dNTP, 0.4 µM of each primer and 0.6 U of AmpliTaq Gold® 360 DNA Polymerase. The PCR profile consisted of initial denaturation in 95 °C for 10 min, then 50 cycles of 95 °C for 30 s, 50 °C for 30 s and 72 °C for 30 s, and finally 72 °C for 7 min. To identify possible mis-incorporated bases caused by post-mortem changes, we PCR amplified and sequenced every region at least twice. As a control, some samples were also extracted twice. Negative controls were included in each extraction and PCR batch. PCR products were purified using Exonuclease I (Thermo Fisher Scientific) and Shrimp Alkaline Phosphatase (Thermo Fisher Scientific). Sequencing reactions were then done using BigDye Terminator v1.1 Cycle Sequencing Kit (Thermo Fisher Scientific), and the products were run on ABI 3730 DNA Analyzer (Applied Biosystems). DNA sequences were edited and assembled into contigs using CodonCode Aligner v4.0.4 (CodonCode Corporation). We also included previously unpublished sequence data from additional modern samples from Lake Saimaa (*N* = 5) and the Baltic Sea (*N* = 2) (for methods, see Valtonen et al. (2012)).

At the Durham University, DNA extraction was carried out following the method described in de Bruyn et al. (2009). The mtDNA control region was amplified in three overlapping fragments of 185-210 bp, from which 490 bp sequence was reconstructed (for primers, see Supplementary Table 2). PCR reactions were performed in a volume of 16 µl containing 1X Qiagen Multiplex PCR Master Mix, 0.2 µM of each primer, and 5 µl of the 5-times diluted extract. The thermal cycling profile was as follows: 95 °C for 15 min, 45 cycles of 94 °C for 30 s, 50 °C for 90 s, and 72 °C for 60 s, followed by a final extension at 72 °C for 10 min. Negative controls were used for all PCR runs. PCR products were purified using QIAquick PCR Purification Kits (Qiagen) according to the manufacturer’s instructions. Purified PCR products were sequenced at the sequencing facility of Durham University. As in Oulu, each sample was sequenced in both directions, and the chromatograms of each sequence were checked by eye.

We obtained the full target region of circa 704 bp from 56 out of 63 museum specimens, 49 of which were fully replicated. A partial target region was obtained from four additional specimens, three of which were fully replicated. We included samples that had unreplicated sequence areas in the analyses, after replacing any unique SNPs in their un-replicated parts with IUPAC ambiguity codes. For example, if the un-replicated part had a thymine (T) in a position in which all other Saimaa ringed seals had a cytosine (C), the T was changed to Y. We additionally included in the analyses five previously unreported modern sequences from Saimaa and two from the Baltic Sea (Supplementary Table 1). All newly reported sequences have been deposited in GenBank under accession numbers XXXXXX-XXXXXX.

**Table 1.**
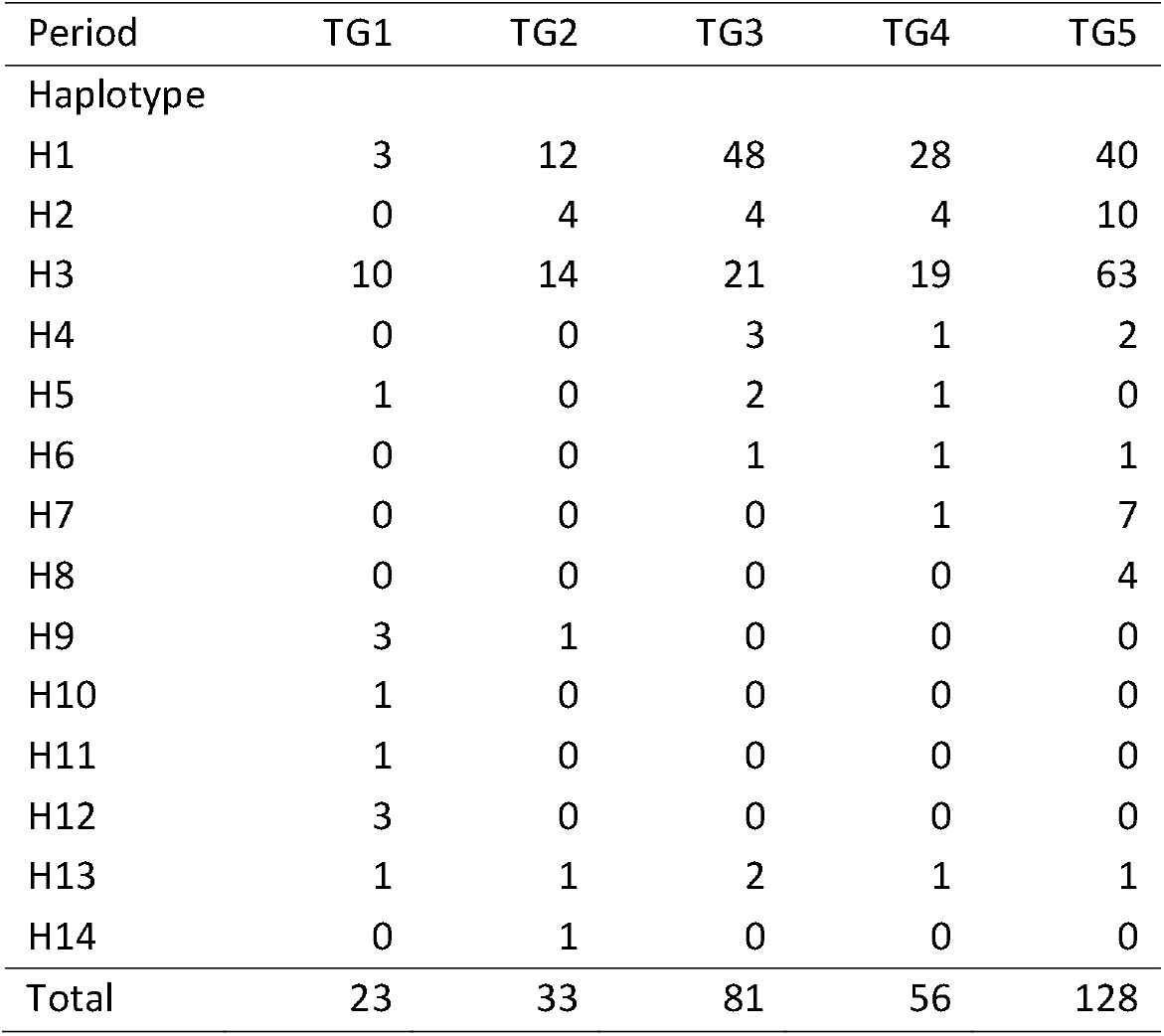
Number of each haplotype observed in each temporal period.

| Period | TG1 | TG2 | TG3 | TG4 | TG5 |
| --- | --- | --- | --- | --- | --- |
| Haplotype |  |  |  |  |  |
| H1 | 3 | 12 | 48 | 28 | 40 |
| H2 | 0 | 4 | 4 | 4 | 10 |
| H3 | 10 | 14 | 21 | 19 | 63 |
| H4 | 0 | 0 | 3 | 1 | 2 |
| H5 | 1 | 0 | 2 | 1 | 0 |
| H6 | 0 | 0 | 1 | 1 | 1 |
| H7 | 0 | 0 | 0 | 1 | 7 |
| H8 | 0 | 0 | 0 | 0 | 4 |
| H9 | 3 | 1 | 0 | 0 | 0 |
| H10 | 1 | 0 | 0 | 0 | 0 |
| H11 | 1 | 0 | 0 | 0 | 0 |
| H12 | 3 | 0 | 0 | 0 | 0 |
| H13 | 1 | 1 | 2 | 1 | 1 |
| H14 | 0 | 1 | 0 | 0 | 0 |
| Total | 23 | 33 | 81 | 56 | 128 |

### 2.2 Haplotype network

A haplotype network for seals from the Baltic area was constructed based on a combined dataset including sequences of museum and modern samples. For the museum samples, only the 56 samples from which the full target region was obtained were included. The modern data included sequences from 208 Saimaa, 21 Baltic and 16 Ladoga ringed seals sampled during the years 1980–2008, 2006– 2009 and 1991–2000, respectively (Valtonen et al., 2012), as well as 57 sequences obtained from Saimaa ringed seal placentas collected from breeding sites during the years 2009–2011 (Valtonen et al., 2015). Sequences were aligned using the FFT-NS-i method within the online version of MAFFT v7 (https://mafft.cbrc.jp/alignment/server/; (Katoh et al., 2002, 2019; Katoh & Standley, 2013)). A total of 24 bases from the beginning and 20 bases from the end of the alignment were excluded to accommodate the seven previously unpublished modern sequences. The final alignment included 663 bp from 358 samples. We first converted the Fasta-formatted alignment file into a TCS input file using FaBox (Villesen, 2007), and then built a haplotype network using TCS (Clement et al., 2000), treating gaps as 5th state. The network was visualized with tcsBU (Múrias Dos Santos et al., 2016).

### 2.3 Temporal genetic variation

To study temporal genetic variation in the Saimaa ringed seal population, the full-length sequences from Saimaa (*N* = 321) were divided into five temporal groups (TG) based on the history of the population outlined in Kunnasranta et al. (2021): TG1 pre–bottleneck 1894–1939 (*N* = 23), TG2 bottleneck 1960–1979 (*N* = 33), TG3 bottleneck 1980–1989 (*N* = 81), TG4 recovery 1990–1999 (*N* = 56), and TG5 increase 2000–2011 (*N* = 128).

Temporal changes in the haplotype frequencies in the Saimaa ringed seal population were visualized as a haplotype network divided into the aforementioned five temporal groups. The figure was drawn manually by editing a network built with TempNet (Prost & Anderson, 2011). We tested for the existence of differences in the spatial distribution of samples across the different temporal groups using chi-square tests. For this, we first excluded the individual from Lake Pielinen, currently uninhabited by seals, as well as nine historical specimens for which the collection site within Lake Saimaa was unknown. To improve concordance to the test’s assumptions, we pooled time periods TG1 and TG2, and combined Kolovesi and Haukivesi into a single area, resulting in a contingency table of 4 time periods by 4 areas. As over 20% of the expected frequencies were still below 5, we conducted a second test using a 5 time periods by 2 areas dataset, in which sampling locations were coded as ‘northern’ (Northern Saimaa, Kolovesi, and Haukivesi basins) or ‘southern’ (Pihlajavesi and Southern Saimaa basins).

As the sample sizes per temporal group are relatively small, the significance of the haplotype frequency differences between the contemporary population (TG5) and earlier temporal groups (TG1–TG4) was tested using a resampling approach. For each haplotype in temporal groups TG1– TG4, *N* observations were drawn with a probability of success equal to the frequency of that particular haplotype in the reference population (TG5) and *N* equal to the sample size in the temporal group in question. For the six haplotypes absent from the reference population TG5, resampling was performed using a frequency of 0.023132 instead of zero, as a haplotype of that frequency would have a 5% chance of not showing up in a sample of *N* = 128. The resampling was repeated 100,000 times per haplotype and temporal group, and the resultant empirical distribution of the expected number of occurrences of each haplotype was recorded and compared with the observed number. The underlying idea is that a nonsignificant haplotype frequency difference may be due to sampling variation, whereas a significant difference could be caused by other factors, such as genetic drift or natural selection. The resampling was performed with an in-house script written for R v3.6.0. by ES (Översti et al., 2019).

### 2.4 Past effective female population size

We studied changes in the past effective female population size of the Saimaa ringed seal by constructing Bayesian skyline plots (Drummond et al., 2005) in BEAST v1.10.4 (Suchard et al., 2018) based on the 321 full-length sequences. The best-fitting substitution model was inferred using model averaging with bModelTest (Bouckaert & Drummond, 2017) in BEAST v2.6.3 (Bouckaert et al., 2019) and analysing the results with Tracer v.1.7.1 (Rambaut et al., 2018). The resulting optimal substitution model was TN93 (Tamura & Nei, 1993) accounting for proportion of invariant sites and rate variation across sites modelled with four gamma categories (Yang, 1994). The data was fitted into two molecular clock models: strict and relaxed uncorrelated lognormal clock. The most suitable clock model was determined with Akaike’s information criteria through MCMC (AICM) (Baele et al., 2012) implemented in Tracer v1.6 (Rambaut et al., 2018). A strict-clock model was preferred over a relaxed clock (AICM = 11.86 with 100 bootstrap replicates) and was used in the subsequent analyses. The sampling years of the sequences were used as a source for external calibration of the molecular clock (i.e. ‘tip-calibration’). We performed three parallel analyses, each with 500 million steps. Every 50,000th step was sampled, and the first ten percent of the steps were discarded as a burn-in. Consistency between runs was evaluated in Tracer, and chains were combined with LogCombiner included in the BEAST software package. Tracer v1.7.2 (Rambaut et al., 2018) was used to confirm that an adequate effective sample size (ESS>200) had been reached for each parameter and for visualization of the Bayesian skyline plot.

Furthermore, although it has been shown that tip-calibration yields more consistent results than internal node calibration (Rieux et al., 2014), on an evolutionary scale, samples in this study can be considered recent. Thus, to assess if the temporal structure of the data used is sufficient to calibrate the molecular clock, we conducted a date-randomization test. We performed twenty random shufflings of tip dates with the R package TIPDATINGBEAST v1.1-0 (Rieux & Khatchikian, 2017) and ran the data-randomized datasets with BEAST v1.10.4 as described above.

### 2.5 Phylogenetic relationships between ringed seal populations

Finally, we studied broader phylogenetic connections among ringed seals from Lake Saimaa and other representative subspecies and populations in the Arctic and Baltic regions. As an addition to the data described above, sequences from Palo (2003) and Martinez-Bakker et al. (2013) were included in this analysis. As for the Baltic region dataset mentioned above, sequences were aligned using the FFT-NS-i method in MAFFT. We used only the sequence region that overlapped in all the datasets, which resulted in an alignment of 348 bp. Next, sequences that had any ambiguous bases in polymorphic sites were removed, bringing the number of sequences down to 551. In order to make the data more manageable for the analysis, sequences were collapsed into 211 unique haplotypes using the DNA to haplotype collapser and converter in FaBox (Villesen, 2007). A Bayesian phylogenetic analysis of the sequence dataset was run in MrBayes v3.2.7a (Ronquist et al., 2012) on the CIPRES Server (Miller et al., 2010). To retain information included in alignment gaps, gaps were recoded to binary presence/absence characters following the method of Simmons & Ochoterena (2000) as implemented in FastGap v1.2 (Borchsenius, 2009). In the run, we used a mixed substitution model with gamma-categorized rate variation for nucleotide sites, while the recoded gaps were treated as a separate partition of variable, gamma-corrected ‘restriction’ sites. Ratepr, statefreqpr, shape, revmat, and tratio were left as default and unlinked across partitions, but the branch length prior was set to unconstrained:Exp(100.0). Two runs with four incrementally heated chains each were run for 10 million generations, while sampling trees every 1,000 generations. Run convergence was assessed in Tracer and, after removing the first 10% of the sampled trees as a burn-in, a midpoint-rooted summary tree showing all compatible groupings was calculated in MrBayes.

## 3 RESULTS

### 3.1 Haplotype network

Our haplotype network for the Baltic area ringed seal mtDNA variation (Fig. 2) shows similar haplotype assemblage as in Valtonen et al. (2012, 2015), but the museum samples carried four additional Saimaa-specific haplotypes (H9, H10, H12 and H14) while lacking four previously observed haplotypes (H4, H6, H7 and H8). In total, the ‘Saimaa clade’ of the network consists of fourteen distinct haplotypes (H1–H14), with one haplotype from the Baltic Sea and the single sequence from Lake Pielinen (H11) being placed in the center of this assemblage. We emphasize that Lake Pielinen has not harbored a permanent ringed seal population in historical times, so the sample from this lake clearly represents a migrant individual that has strayed to Lake Pielinen from Lake Saimaa via the 67 km long River Pielisjoki. As in the results of Valtonen et al. (2012), the network shows all the Lake Saimaa haplotypes clustering relatively close to each other, with two haplotypes (H1 and H3) dominating through time. Haplotype H14 encountered in Lake Saimaa has a 64-bp deletion but also additional SNPs, which explain its outlier position in the network.

**Figure 2.**
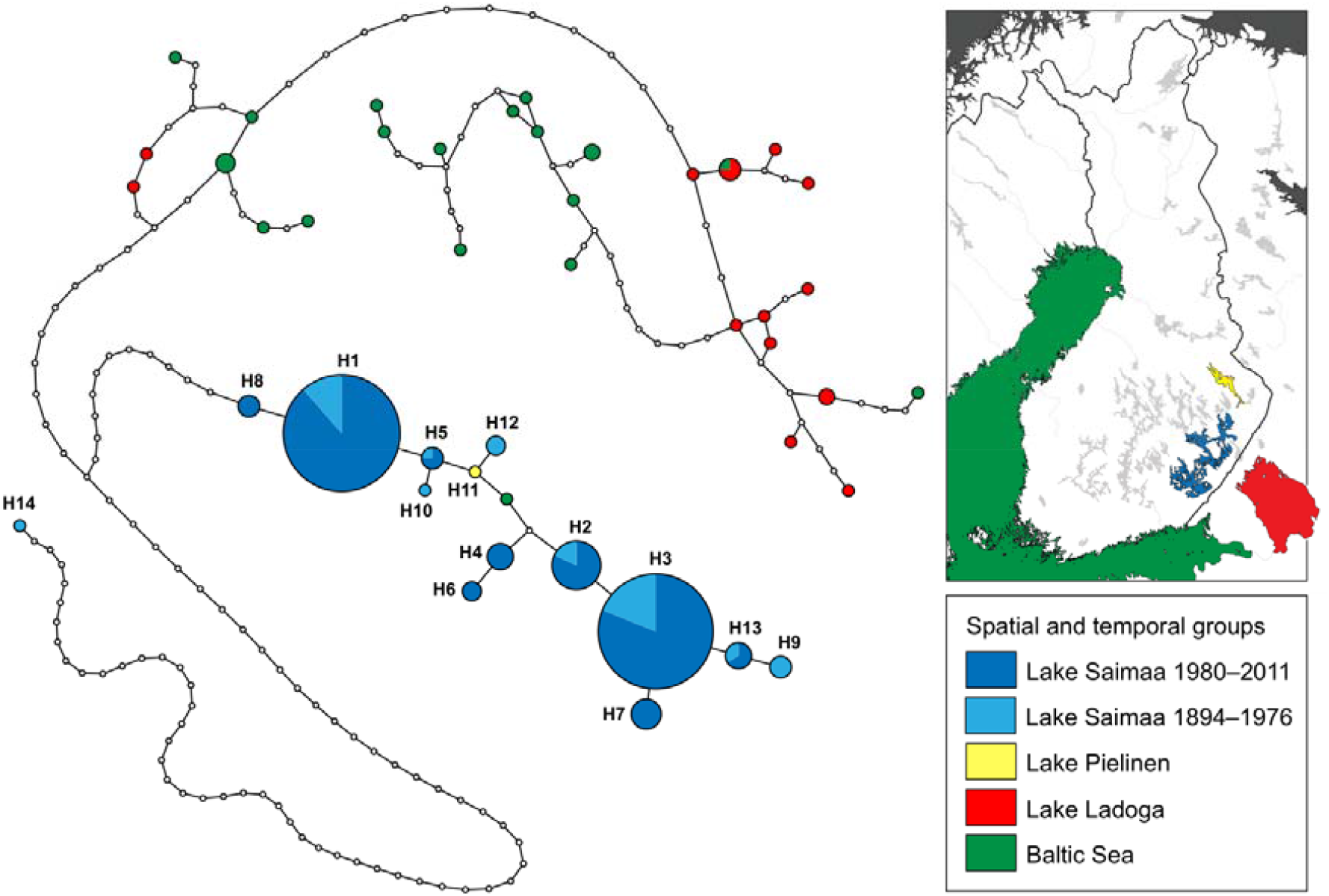
Haplotype network of ringed seals in the Baltic Sea and lakes Saimaa, Pielinen, and Ladoga. The sizes of the circles correspond to the number of sampled individuals with the haplotype, section colors denote sampling site and collection period within Lake Saimaa (see map and legend in inset). Small white circles denote unobserved haplotypes. Note that haplotype H14 is present in a single individual that has a 64-bp deletion (each bp in the deletion is counted as one change) as well as unique substitutions.

### 3.2 Temporal genetic variation

Division of the samples into five temporal groups (TG1–TG5) allowed exploring changes through time in the Lake Saimaa mtDNA haplotype pool. The temporal haplotype network (Fig. 3) and the haplotype table (Table 1) show substantial fluctuation in haplotype frequencies through time. The most common haplotype today, H3, seems to have been dominant also during the two first periods (TG1: 1894–1939 and TG2: 1960–1979). The currently second most common haplotype, H1, however, seems to have become common only after TG1: 1894–1939, and it was even more common than H3 during TG3: 1980–1989 and TG4: 1990–1999. The third most common haplotype (H2) is observed only from period TG2: 1960–1979 onwards, with a slight increase in frequency towards the Present.

**Figure 3.**
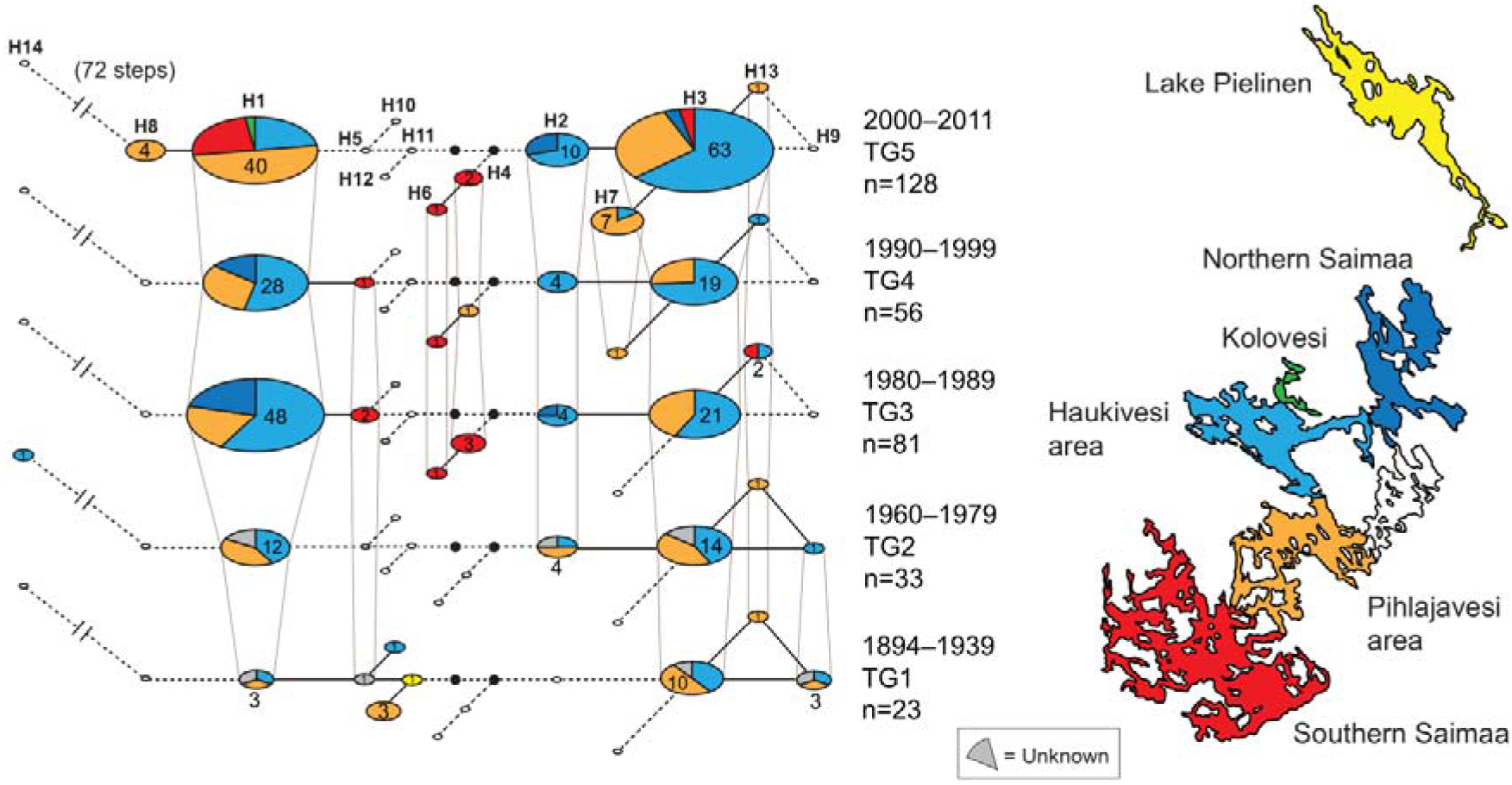
Haplotype network of Saimaa ringed seals during five different time intervals spanning periods from 1894 to the Present. In each network, circle size is proportional to haplotype frequency, and sector colors denote sampling areas as shown in the map to the right of the networks.

Despite the relatively small temporal group-specific sample sizes, our resampling tests (Fig. 4) showed that many of the observed differences in haplotype frequencies are unlikely to be caused by mere sampling variation: five haplotypes in TG1–TG4 showed a nominally significant difference from TG5 within at least one time period (seven significant differences in all). While this result is not corrected for multiple testing (i.e., for the 4 × 14 = 56 haplotype-wise tests), only 2.8 significant results would be expected by chance at a significance level of α = 0.05 in 56 independent tests, and the probability of obtaining seven or more significant results by chance would be 0.021.

**Figure 4.**
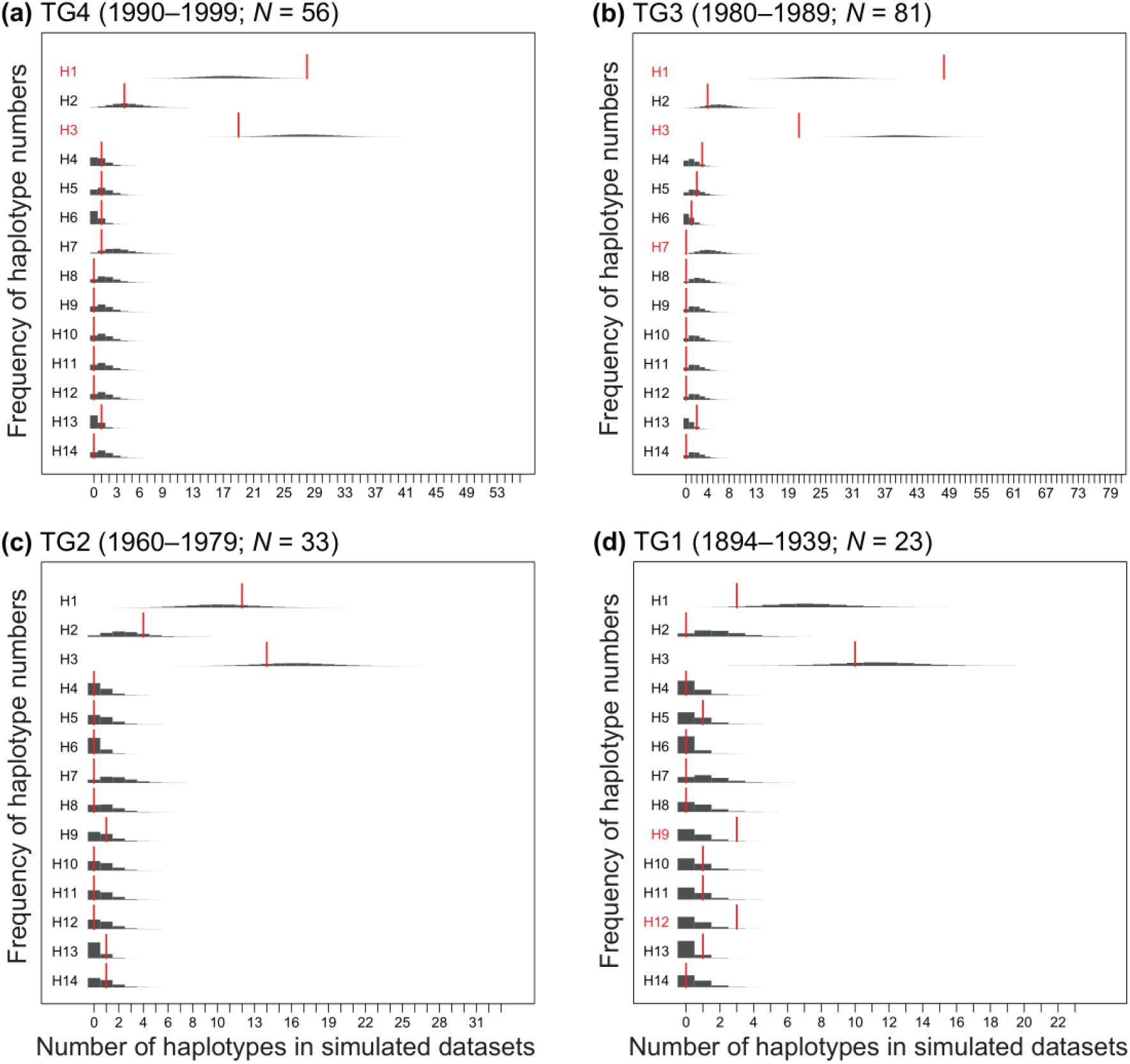
Haplotype frequency differences between the modern population and earlier temporal groups (TG1–TG4). Dark grey bars depict the expected distribution of the number of observations per haplotype in each temporal group based on 100,000 rounds of resampling from the modern distribution (TG5, *N* = 128). Red bars indicate the observed numbers within each temporal group, and red font denotes haplotypes with a nominally significant frequency difference as compared the modern population (*i*.*e*., with an observed number of occurrences outside 95% of the simulated probability mass).

Taking into account also the Lake Pielinen specimen, eight different haplotypes were observed in the earliest time interval TG1 (1894–1939). Three of these (H10, H11 and H12) were not observed in later time. The resampling analysis suggested that the overrepresentation of H9 and H12 in this time interval cannot be explained by small sample size alone (Fig. 4). H12 is only observed in TG1, with a frequency of ≈ 13%. The fact that three unique haplotypes were observed only in the smallest sample of the oldest time period shows that some variation has been permanently lost and also suggests that some of the past variation remains unsampled. In TG2 (1960–1979), only six haplotypes were observed despite a larger sample size than in TG1, suggesting substantial loss of haplotype diversity between 1894 and 1960. TG3 (1980–1989) demonstrates a step towards the modern haplotype distribution, with H1 and H3 dominating. Nevertheless, compared to modern times, H1 is nominally significantly overrepresented and H3 underrepresented both in the 1980s and 1990s samples (TG3 and TG4). Furthermore, H7, which has a frequency of ≈ 5% in the modern TG5 (2000–2011), appears for the first time in TG4 (1990–1999). Haplotype H8 is only observed in TG5 (2000–2011) despite the larger summed sample size of the earlier time periods (*N* = 128 vs. *N* = 193).

Our tests for differential sampling across time periods indicated differences in the 4 temporal groups by 4 areas dataset (χ2 = 12.7, df = 9, *P* = 0.0005), but in this case the assumptions of the test are slightly violated (over 20% of the expected frequencies were below 5). The test assumptions were, however, met by the 5 temporal groups by 2 areas dataset, which also resulted in statistically significant differences in sampling (*χ*2 = 12.744, df = 4, *P* = 0.013).

### 3.3 Past effective female population size

The Bayesian skyline plot depicting the past effective population size of Saimaa ringed seal females (Fig. 5) suggests that the population has slowly decreased in size at least since the beginning of the 12^th^ century, but with an abrupt escalation of the population decline from the middle of the 19th century onwards. In our date-randomization test, the 95 % highest posterior densities for the Clock.rate and TreeModel.rootHeight parameters overlapped somewhat between the actual and randomized datasets (Supplementary Fig. S1).

**Figure 5.**
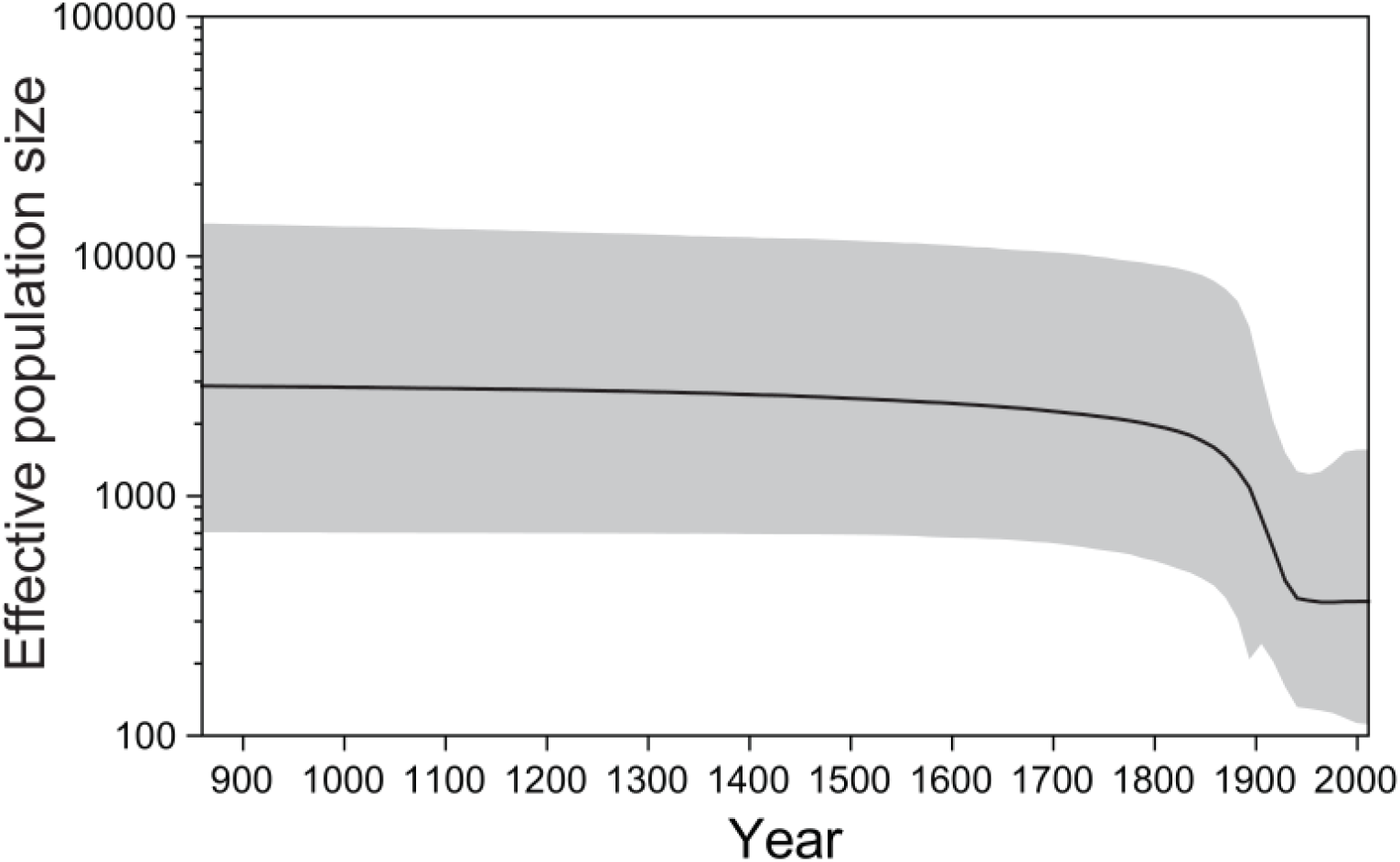
Bayesian skyline plot depicting the estimated historical effective population size of Saimaa ringed seal females on a logarithmic scale. Calendar years are shown on the X-axis. The thick line shows the median and the shaded area represents the 95% highest posterior density interval.

### 3.4 Phylogenetic relationships among ringed seal populations

In our ringed seal mtDNA phylogeny (Fig. 6), the Saimaa ringed seal clade is unexpectedly nested within the Arctic, and more specifically North American, ringed seal diversity rather than the Baltic ringed seal diversity. The ringed seals from the White Sea, which are geographically the closest Arctic population to Saimaa, are however not closely related to Saimaa. This is also the case for the geographically adjacent Ladoga ringed seal population, as individuals from Lake Ladoga are widely scattered throughout the phylogeny. Interestingly, the aforementioned Saimaa ringed seal individual 5688 (haplotype H14) with the 64-bp deletion is grouped with 1.0 posterior support together with two North American individuals that share the same deletion as well as SNPs. Five Baltic ringed seal haplotypes are nested inside the Saimaa ringed seal clade.

**Figure 6.**
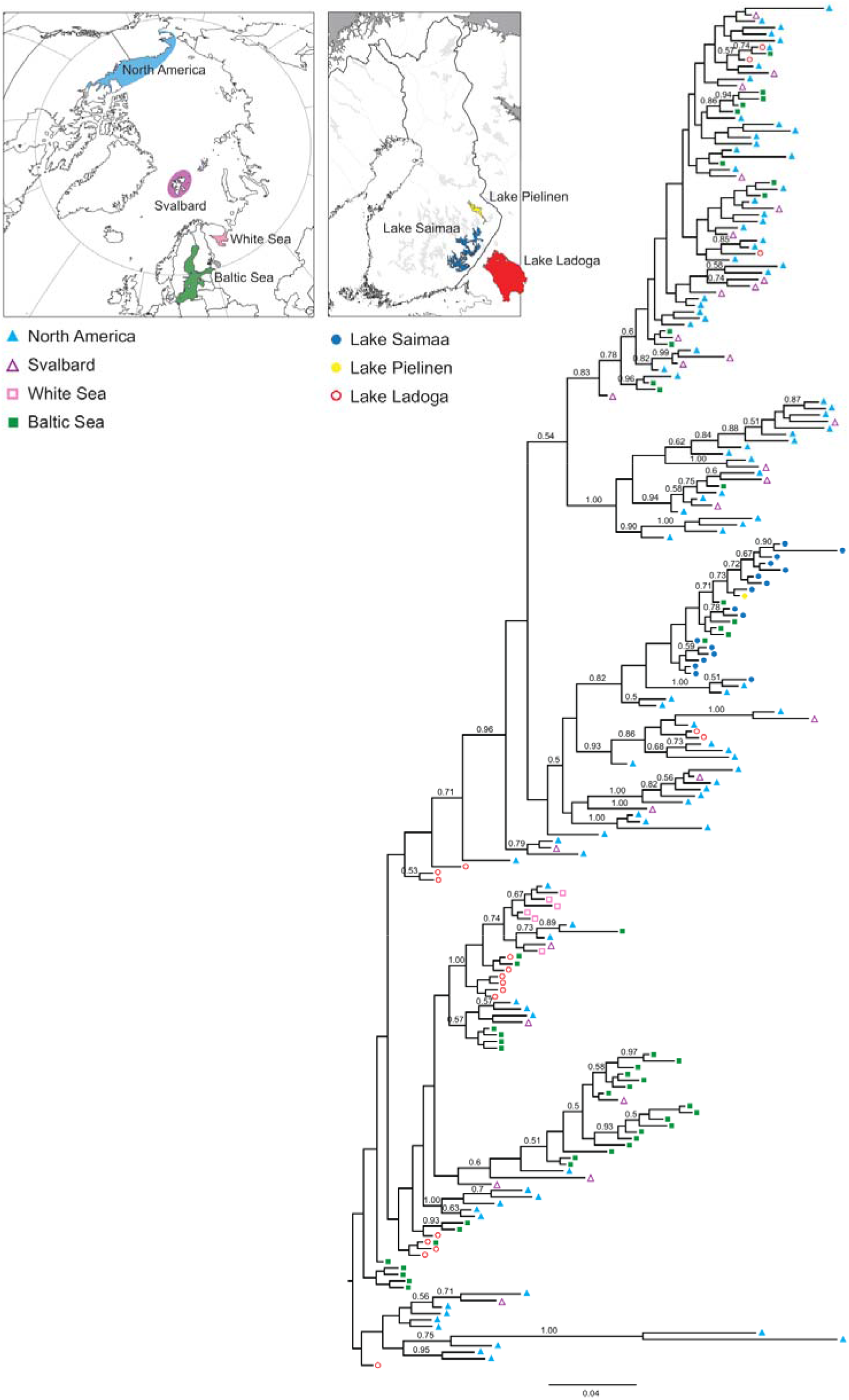
Bayesian phylogenetic tree of 211 unique mitochondrial control-region haplotypes found in freshwater, Baltic and Arctic ringed seal populations. Symbols and colors at the tips of the tree indicate the areas where each haplotype is present (see inset map and legend). Numbers above branches are posterior probabilities, only values over 0.50 are shown.

## 4 DISCUSSION

Improvements in molecular-genetic methods have increasingly made museum collections an invaluable source for gaining insights into the past genetic composition of animal and plant species (Burrell et al., 2015; Nakahama, 2021; Raxworthy & Smith, 2021). Museum material allow inferences of long-term changes in the frequencies of genetic variants (Bi et al., 2019), and can be used to improve estimates of demographic trajectories (Hung et al., 2014). Especially for endangered species, genotyping museum samples can allow direct demonstration of loss of genetic diversity over time due to diminished population size (Dussex et al., 2021; van der Valk et al., 2019; von Seth et al., 2021). Here, we sequenced mitochondrial control-region sequences from 60 museum-preserved samples of the endangered Saimaa ringed seal, and then combined these data with previously published contemporary sequences from Lake Saimaa. We used this combined dataset spanning 117 years to infer the genetic impacts of the 20^th^-century population bottleneck, caused mainly by relentless persecution of Saimaa ringed seals during the first half of the century. We also combined our Saimaa dataset with existing data from other Holarctic ringed seal subspecies, in order to investigate the origin of the lake-endemic Saimaa ringed seal population.

### 4.1 The genetic signature of the 20^th^ -century bottleneck

The Saimaa ringed seal ranks among the genetically least diverse animal subspecies on the Earth, both in terms of mitochondrial (Martinez-Bakker et al., 2013; Nyman et al., 2014; Palo, 2003; Valtonen et al., 2012, 2014) and nuclear (Martinez-Bakker et al., 2013; Nyman et al., 2014; Palo, 2003; Palo et al., 2003; Stoffel et al., 2018; Valtonen et al., 2014, 2015) genetic diversity. The genetic uniformity of the subspecies is even more striking when considering the fact that ringed seals in general tend to be highly diverse (Martinez-Bakker et al., 2013; O’Corry-Crowe, 2008; Olsen et al., 2011; Palo et al., 2001). As estimated by Nyman et al. (2014), Palo et al. (2003) and Valtonen et al. (2012), the Saimaa ringed seal has lost 33.9% and 89.4% of its mitochondrial haplotype and nucleotide diversity respectively, as well as 55-69% of its overall heterozygosity in relation to its relatives in the Arctic Ocean and the Baltic Sea.

As such, the anthropogenic population collapse of the Saimaa ringed seal in the 20^th^ century is well known and thoroughly documented (Hyvärinen et al., 1998; Kokko et al., 1998; Ranta et al., 1996; Sipilä et al., 1990). However, whether this bottleneck fully explains the extremely low genetic diversity of the population is less evident. The Saimaa ringed seal is long-lived and has an estimated generation time of 11 years (Palo et al., 2001). Hence, Nyman et al. (2014), Palo et al. (2003) and Valtonen et al. (2012) inferred that the main portion of diversity loss occurred well prior to the recent bottleneck, *i*.*e*., during the long post-glacial isolation of the Saimaa ringed seal. Nyman et al. (2014) used Approximate Bayesian Computation inference based on combined nuclear and mitochondrial data and postulated that incorporating the recent bottleneck does not improve model fit in relation to a demographic model that includes only a bottleneck during colonization of the Lake Saimaa system nearly 10,000 years ago.

However, it should be emphasized that abrupt or slow post-glacial genetic erosion does not preclude the possibility that the recent bottleneck would have led to further marked loss of diversity. Indeed, our Bayesian skyline plot (Fig. 5) showed a sharp reduction in female effective population size towards the Recent. The commencement of the dip in the estimated trajectory closely parallels the recent population collapse, which started in the early 19th century and intensified during the first half of the 20^th^ century, largely as a result of a growing human population and increased access to modern firearms in the region. In this respect, our results are also consistent with the recent results of Stoffel et al. (2018) and Peart et al. (2020), who found statistically significant signatures of the recent bottleneck based on nuclear microsatellites and ddRAD-based SNP’s, respectively. The effects of the still-continuing population recovery, which started in the 1980s, were not manifested in our most recent temporal samples of 2011.

The patterns we observed in the Bayesian skyline plot need to be interpreted with caution, as the 95% posterior density interval of the trajectory is wide, and population substructure and unbalanced sampling may confound skyline analyses (Heller et al., 2013). Moreover, in our date-randomization test, the 95% highest posterior densities for the clockRate and TreeHeight parameters overlapped somewhat between the actual and randomized datasets (Supplementary Fig. S1). This implicates that the temporal timeframe of the sampled sequences might not be wide enough to produce entirely reliable estimates of evolutionary rate and time. Future analyses should aim at strengthening the temporal signal by including older samples, longer sequences, and/or internal node calibrations, as suggested by Rieux & Khatchikian (2017).

### 4.2. Fluctuations in haplotype frequencies and recent loss of diversity

We observed substantial fluctuation in mtDNA haplotype frequencies through time in the Saimaa ringed seal population. While Valtonen et al. (2012, 2014) demonstrated significant differentiation in haplotype frequencies between the years 1980 and 2008, our analyses incorporating museum material extend the pattern nearly a hundred years back in time. As shown by our temporal networks (Fig. 3) and resampling analyses (Fig. 4), several haplotypes have probably been completely lost, while some haplotypes that are dominant today have only recently increased in frequency. The observed differences in haplotype frequencies in the different temporal groups are likely real and not caused by sampling bias. The observation of three unique haplotypes in the sample of 23 individuals that lived between 1894–1939, but not found among the nearly 300 individuals from later times, provides concrete evidence for diversity loss. Given the small sample sizes in the oldest temporal groups, it is also likely that much of the past variation remains unsampled. The observed haplotype loss and drastic frequency changes suggest that, during the last 125 years, the reduction in effective female population size in the Saimaa ringed seal has been dramatic. In this, the fate of the Saimaa ringed seal resembles another severely persecuted mammal, the grey wolf (*Canis lupus*). The wolves in the past have harbored high mtDNA diversity that has collapsed in the turn of the 20^th^ century especially in Western Europe (Dufresnes et al., 2018).

When interpreting the results above, a potential complication arises because of the previously documented genetic substructure within Lake Saimaa (Valtonen et al., 2014), which could bias inferences if the main regions of the lake are differentially represented in the temporal samples. Statistically significant differential sampling indeed seemed to be present in the dataset. The fact that samples from Southern Saimaa are only present in the three most recent temporal groups (TG3, TG4 and TG5) might explain the absence of some Southern Saimaa-specific haplotypes in the earlier time intervals and would at the same time bias estimated diversity of the most recent time periods upwards. However, regionally biased sampling does not seem to explain the loss of haplotypes H9, H10, H12 and H14 in the most recent time intervals, as these haplotypes were observed in lake regions that are well represented in temporal groups TG3–TG5.

### 4.3 Origin of the Saimaa ringed seal

According to the commonly held view, the Saimaa ringed seal population originated from Baltic ringed seals trapped in an inland lake system that gradually emerged due to post-glacial land uplift around 9,000 years ago (Hyvärinen & Nieminen, 1990; Nyman et al., 2014). However, the mtDNA phylogeny presented here (Fig. 6) suggests that the modern Baltic ringed seal population is a poor proxy for the ancestral Saimaa population. All the Saimaa haplotypes form a cluster that is not nested within the Baltic haplotypes. In fact, none of the geographically adjacent populations in the Baltic Sea, Lake Ladoga, and the White Sea are close to the Saimaa ringed seal cluster. Rather, the Saimaa ringed seal clade is nested within the Arctic, especially North American, ringed seal diversity, which suggests that the history of the Saimaa ringed seal population is more complex than previously thought. The fact that some Baltic ringed seal haplotypes are nested inside the Saimaa ringed seal mtDNA clade (Figs. 2 and 6) might conceivably suggest returning gene flow from Lake Saimaa into the Baltic population. However, the current outlet of Lake Saimaa is River Vuoksi, which runs into Lake Ladoga, from which the route to the Baltic Sea continues via River Neva. The absence of Saimaa-clade haplotypes from Lake Ladoga therefore suggests that their presence in the Baltic Sea reflects incomplete lineage sorting.

As already noted by Palo (2003) and Valtonen et al. (2012), particularly problematic for the post-glacial ‘Out of Baltic’ scenario for the Saimaa ringed seal is the presence of multiple closely related, but still diverse, haplotypes in Lake Saimaa. It is unlikely that a random sampling (through lineage sorting either during the initial colonization phase or later) from the Baltic haplotype pool would retain such a closely related set of haplotypes. The presence of a single mtDNA clade within Lake Saimaa could theoretically be explained by an extended colonization bottleneck (Nyman et al., 2014) and differentiation of a few founding lineages during the isolation. The current diversity within Lake Saimaa is, however, too high to have emerged post-glacially. As pointed out by Palo (2003), the accumulation of the diversity observed in the Saimaa clade would require at least 95,000 years (c. 8,600 generations), or Saimaa-specific mutation rates that are roughly 10x faster than normally estimated for mammalian mtDNA.

The observed pattern could have emerged if the Baltic Sea area has experienced multiple colonization waves of ringed seals after the last glacial period (Schmölcke, 2008), so that the Saimaa population represents survivors of an early wave, and the extant Baltic and Ladoga populations derive from a later wave that replaced the earlier Baltic population. Based on ancient specimens, Bro-Jørgensen (2021) inferred that such a replacement has happened in the Baltic Sea grey seal (*Halichoerus grypus* (2014). However, according to Ukkonen et al. (2014), there is no paleontological evidence suggesting that the Baltic ringed seal population would have experienced extinctions and recolonizations after the initial immigration into the basin after the last glacial period, though additional immigration from the Arctic may have occurred.

On the other hand, the pattern could be explained if Saimaa ringed seals originated from an ancestral population that inhabited proglacial lake systems situated on the edge of the Fennoscandian Ice Sheet. Current models support the existence of such refugia other aquatic species, such as the Atlantic salmon (*Salmo salar* L.) (Asplund et al., 2004; Nilsson et al., 2001; Tonteri et al., 2005, 2007), grayling (*Thymallus thymallus*) (Koskinen et al., 2000), and European perch (*Perca fluviatilis*) (Nesbø et al., 1999).

### 4.4 Conclusions

The use of museum specimens allowed us to directly investigate genetic patterns in the endangered Saimaa ringed seal population through more than a hundred years. Although erosion of the initial genetic diversity has continued throughout the isolation, we observed 20^th^ century loss of haplotypes and relatively drastic fluctuations in haplotype frequencies, demonstrating a genetic effect of the human-induced population collapse. Combining newly generated and already published data from multiple ringed seal populations additionally allowed us to investigate broad phylogeographic patterns in ringed seals. In apparent contrast with the traditionally held view of the Baltic origin of the population, the Saimaa ringed seal mtDNA variation shows enigmatic affinities to North American ringed seals. These results add to the growing body of evidence which calls for a re-evaluation of the deep history of the Saimaa ringed seal population. Future data on still-unsampled populations, for example in the Arctic Ocean, as well as genomic data and ancient DNA could provide keys to understanding the origin and demographic history of the Saimaa ringed seal. Lake Saimaa may harbor a ringed seal population that is even more unique than previously thought.

## Acknowledgements

This work was carried out with the support of the Centre for Material Analysis, University of Oulu, Finland. The authors wish to acknowledge CSC – IT Center for Science, Finland, for computational resources and the Finnish Museum of Natural History for access to their collections. MTH acknowledges funding from the Emil Aaltonen Foundation and the University of Oulu Scholarship Foundation. MV was funded by the Maj and Tor Nessling Foundation and Jane and Aatos Erkko Foundation. MP was supported by Marie Skłodowska Curie Intra-European Fellowship from the European Commission (PIEF-GA-2009-235978) and the Polish National Agency for Academic Exchange (NAWA; Polish Returns Fellowship PPN/PPO/2018/1/00037). ES acknowledges funding from the Jenny and Antti Wihuri Foundation and the Ella and Georg Ehrnrooth Foundation. The Raija and Ossi Tuuliainen Foundation, Kuopio Naturalists’ Society and Nestori Foundation are also acknowledged for funding the research. We like to thank Jaakko Lumme and Laura Kvist from the University Oulu and Love Dalén from the Centre for Palaeogenetics for discussion. We further thank Pirkko Ukkonen for her help with the material collection from museum specimens.

## DATA AVAILABILITY STATEMENT

The sequences reported in this study have been deposited in GenBank under accession numbers X. The script used for haplotype resampling is deposited in X.

## Supplementary Material

**Supplementary Table S1**. Studied ringed seal specimens sorted by known/approximate year of death. The table shows sequence ID, catalog ID, subspecies status, whether the full target sequence was obtained, year of death, location, sex, source collection and type of sample/tissue.

**Supplementary Table S2**. Primers used to amplify the mtDNA control region in ringed seals.

**Supplementary Figure S1**. Results from date-randomization test for the (a) Clock.rate and (b) TreeModel.rootHeight parameters. For both parameters, 95% highest posterior densities (HPD) for the true data (red) and for twenty date-randomized datasets (black) are presented on a logarithmic scale.

## Author contributions

TN, MK, JUP, JA, MV, MR, MTH, MP, ARH conceived the idea; JUP, MP, JA performed sampling; MTH, JH, MP performed the laboratory work; MTH, TN, JUP, MV, SÖ, ES, JA conducted the data analyses; TN, MTH, ES prepared the figures; JA, MR supervised the work; MV, TN, MK, JUP, JA, MR, MTH, MP acquired funding; JUP, TN, MTH, MV, MP, SÖ, ES, JA wrote the original draft; all authors except MR reviewed and commented the manuscript.

