## Supplementary figures and images for "Museum specimens of a landlocked pinniped reveal recent loss of genetic diversity and unexpected population connections"

### Supplementary Fig. S1

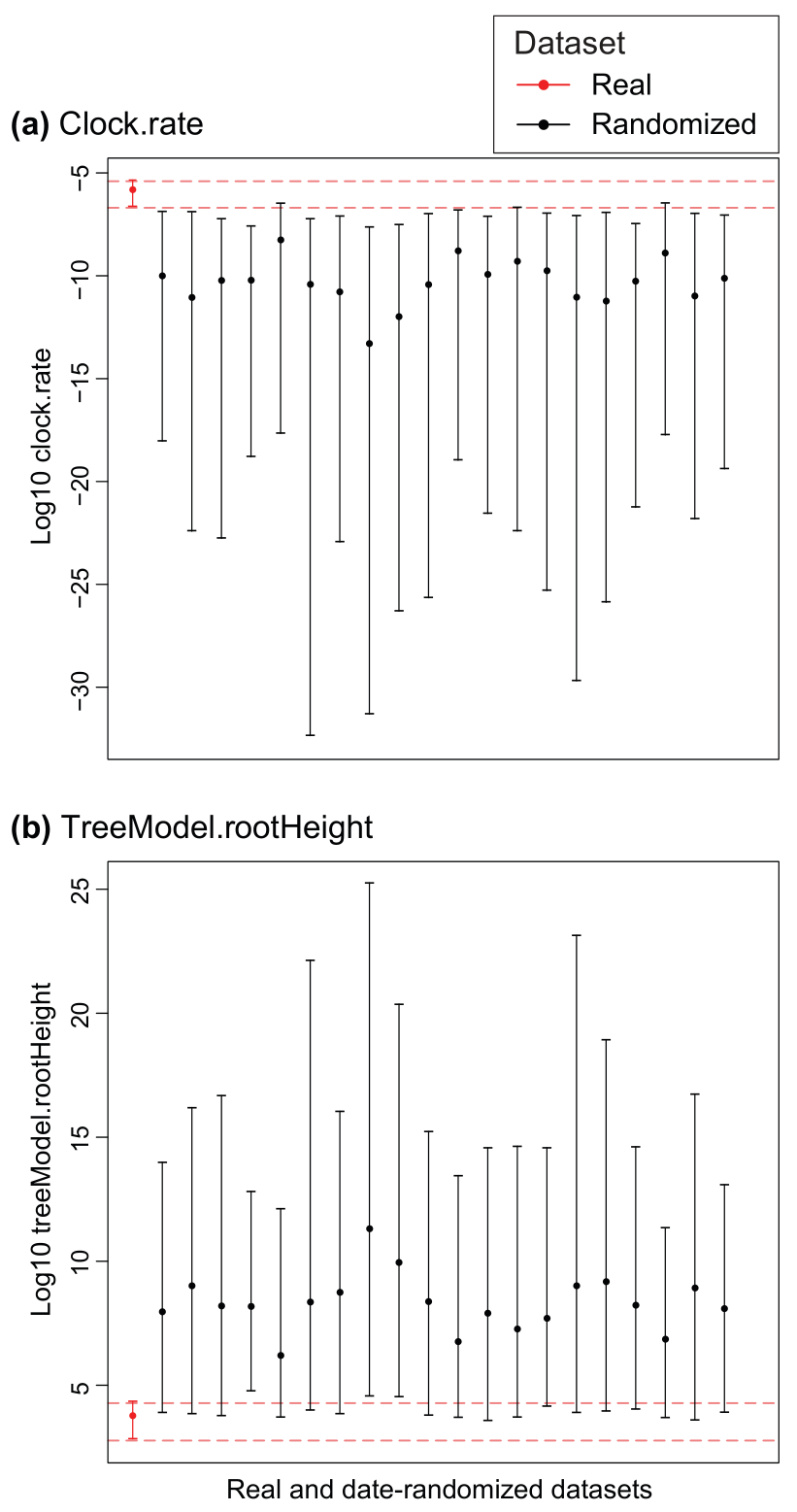
