## Supplementary Table S1 for "Museum specimens of a landlocked pinniped reveal recent loss of genetic diversity and unexpected population connections"

Supplementary Table 1. Studied ringed seal specimens sorted by known/approximate year of death.

| **Sequence ID** | **Catalog ID** | **Subspecies** | **Target region obtained** | **Year of death** | **Location** | **Sex** | **Source**  **collection** | **Type of sample/tissue** |
| --- | --- | --- | --- | --- | --- | --- | --- | --- |
| 2296 | KN 2296 | *P. h. saimensis* | Full | before 1899 | Finland: Lake Saimaa | U | Luomus | Canine |
| 17 | KN 2300(17) | *P. h. saimensis* | Full | before 1899 | Finland: Lake Saimaa | U | Luomus | Two postcanine teeth |
| M426 | KN 2297 | *P. h. saimensis* | Full | before 1899 | Finland: Lake Saimaa | U | Luomus | Bone |
| 19 | KN 2301(19) | *P. h. saimensis* | Full | 1894 | Finland: Lake Saimaa, Haukivesi | U | Luomus | Canine |
| M422 | KN 1488 | *P. h. saimensis* | Full | 1894 | Finland: Lake Saimaa, Rantasalmi | U | Luomus | Bone |
| M357 | 631191 | *P. h. saimensis* | Partial | 1906 | Finland: Lake Saimaa, Olofsborg | U | ? | Bone |
| 1663 | KN 1663 | *P. h. saimensis* | Full | before 1913 | Finland: Lake Saimaa, Enonkoski | M | Luomus | Digital bone and claw |
| 2576 | KN 2576 | *P. h. saimensis* | Full | 1913 | Finland: Lake Saimaa, Pielisjärvi | F | Luomus | Digital bone and claw |
| 2577 | KN 2577 | *P. h. saimensis* | Full | 1913 | Finland: Lake Saimaa, Kerimäki | U | Luomus | Digital bone, claw and skin |
| 3664 | KN 3664 | *P. h. saimensis* | Full | 1913 | Finland: Lake Saimaa, Pihlajavesi | U | Luomus | Two postcanine teeth |
| 6903 | KN 6903 | *P. h. saimensis* | Full | 1913 | Finland: Lake Saimaa, Puruvesi | U | Luomus | Digital bone and claw |
| M27 | KN 2295 | *P. h. saimensis* | Partial | 1913 | Finland: Lake Saimaa, Puruvesi | U | Luomus | Bone |
| 2569 | KN 2569 | *P. h. saimensis* | Full | 1914 | Finland: Lake Saimaa, Enonkoski | F | Luomus | Digital bone and claw |
| 2571 | KN 2571 | *P. h. saimensis* | Full | 1914 | Finland: Lake Saimaa, Enonkoski | M | Luomus | Digital bone and claw |
| 2574 | KN 2574 | *P. h. saimensis* | Full | 1914 | Finland: Lake Saimaa, Enonkoski | F | Luomus | Digital bone and claw |
| 6904 | KN 6904 | *P. h. saimensis* | Full | 1914 | Finland: Lake Saimaa, Enonkoski | M | Luomus | Digital bone, claw and skin |
| 2292 | KN 2292 | *P. h. saimensis* | Full | 1915 | Finland: Lake Saimaa, Kerimäki | U | Luomus | Postcanine tooth |
| 2293 | KN 2293 | *P. h. saimensis* | Full | 1915 | Finland: Lake Saimaa, Punkaharju | U | Luomus | Postcanine tooth |
| 6926 | KN 6926 | *P. h. saimensis* | Full | 1922 | Finland: Lake Saimaa, Puruvesi | M | Luomus | Coccygeal vertebra |
| 6925 | KN 6925 | *P. h. saimensis* | Full | 1923 | Finland: Lake Saimaa, Puruvesi | M | Luomus | Digital bone |
| M350 | 612604 | *P. h. saimensis* | Partial | 1924 | Finland: Lake Saimaa, Punkasalmi | U | Luomus | Bone |
| 6929 | KN 6929 | *P. h. saimensis* | Full | 1925 | Finland: Lake Saimaa, Puruvesi | U | Luomus | Claw |
| 6928 | KN 6928 | *P. h. saimensis* | Full | 1926 | Finland: Lake Saimaa, Puruvesi | U | Luomus | Digital bone and claw |
| 6930 | KN 6930 | *P. h. saimensis* | Full | 1926 | Finland: Lake Saimaa, Puruvesi | U | Luomus | Digital bone and claw |
| 6933 | KN 6933 | *P. h. saimensis* | Full | 1926 | Finland: Lake Saimaa, Puruvesi | U | Luomus | Digital bone and claw |
| M416 | KN 46714 | *P. h. saimensis* | Partial | 1926 | Finland: Lake Saimaa, Puruvesi | U | Luomus | Bone |
| M417 | KN 46802 | *P. h. saimensis* | Full | 1939 | Finland: Lake Saimaa, died at Helsinki Zoo | M | Luomus | Bone |
| 5652 | KN 5652 | *P. h. saimensis* | Full | 1962 | Finland: Lake Saimaa, Haapavesi | M | Luomus | Digital bone |
| 5653 | KN 5653 | *P. h. saimensis* | Full | 1962 | Finland: Lake Saimaa, Sääminki | M | Luomus | Bone |
| 5655 | KN 5655 | *P. h. saimensis* | Full | 1962 | Finland: Lake Saimaa, Pihlajavesi | M | Luomus | Digital bone |
| 5689 | KN 5689 | *P. h. saimensis* | Full | 1964 | Finland: Lake Saimaa, Sääminki, Paatisen luoto | F | Luomus | Postcanine tooth and digital bone |
| 5691 | KN 5691 | *P. h. saimensis* | Full | 1964 | Finland: Lake Saimaa, Sääminki | M | Luomus | Postcanine tooth and digital bone |
| 5692 | KN 5692 | *P. h. saimensis* | Full | 1964 | Finland: Lake Saimaa, Sääminki, Teerivesi | M | Luomus | Claw |
| 5695 | KN 5695 | *P. h. saimensis* | Full | 1964 | Finland: Lake Saimaa, Rantasalmi | F | Luomus | Digital bone and claw |
| 5700 | KN 5700 | *P. h. saimensis* | Full | 1964 | Finland: Lake Saimaa, Tuohiselkä | M | Luomus | Canine, incisor and digital bone |
| 5690 | KN 5690 | *P. h. saimensis* | Full | 1965 | Finland: Lake Saimaa, Rantasalmi | M | Luomus | Canine |
| 5693 | KN 5693 | *P. h. saimensis* | Full | 1965 | Finland: Lake Saimaa, Sulkava | M | Luomus | Fragment of postcanine tooth, digital bone and claw |
| 5694 | KN 5694 | *P. h. saimensis* | Full | 1965 | Finland: Lake Saimaa, Pihlajavesi | F | Luomus | Digital bone and claw |
| 5696 | KN 5696 | *P. h. saimensis* | Full | 1965 | Finland: Lake Saimaa, Rantasalmi | F | Luomus | Fragment of upper jaw with two postcanine teeth |
| 5697 | KN 5697 | *P. h. saimensis* | Full | 1965 | Finland: Lake Saimaa, Pyyvesi | M | Luomus | Incisor and digital bone |
| 5698 | KN 5698 | *P. h. saimensis* | Full | 1965 | Finland: Lake Saimaa, Haukivesi | M | Luomus | Incisor and digital bone |
| 5703 | KN 5703 | *P. h. saimensis* | Full | 1965 | Finland: Lake Saimaa, Pihlajavesi | F | Luomus | Digital bone |
| 5704 | KN 5704 | *P. h. saimensis* | Full | 1965 | Finland: Lake Saimaa, Pihlajavesi | M | Luomus | Digital bone |
| 5688 | KN 5688 | *P. h. saimensis* | Full | 1966 | Finland: Lake Saimaa, Haapavesi | F | Luomus | Postcanine tooth and digital bone |
| 5701 | KN 5701 | *P. h. saimensis* | Full | 1966 | Finland: Lake Saimaa, Haukivesi | F | Luomus | Canine |
| 5687 | KN 5687 | *P. h. saimensis* | Full | 1968 | Finland: Lake Saimaa, Väistönselkä | F | Luomus | Digital bone |
| 6097 | KN 6097 | *P. h. saimensis* | Full | 1970 | Finland: Lake Saimaa, Oravivesi | F | Luomus | Digital bone |
| 6133 | KN 6133 | *P. h. saimensis* | Full | 1970 | Finland: Lake Saimaa, Oravivesi | U | Luomus | Digital bone and claw |
| 6134 | KN 6134 | *P. h. saimensis* | Full | 1970 | Finland: Lake Saimaa, Väistönselkä | F | Luomus | Digital bone and claw |
| 6227 | KN 6227 | *P. h. saimensis* | Full | 1970 | Finland: Lake Saimaa, Pihlajavesi | U | Luomus | Digital bone and claw |
| 6465 | KN 6465 | *P. h. saimensis* | Full | 1970 | Finland: Lake Saimaa, Pihlajavesi | U | Luomus | Postcanine tooth and digital bone |
| 6287 | KN 6287 | *P. h. saimensis* | Full | 1971 | Finland: Lake Saimaa, Haukivesi | M | Luomus | Digital bone and claw |
| 6288 | KN 6288 | *P. h. saimensis* | Full | 1971 | Finland: Lake Saimaa, Haukivesi | F | Luomus | Digital bone and claw |
| 6289 | KN 6289 | *P. h. saimensis* | Full | 1971 | Finland: Lake Saimaa, Pihlajavesi | F | Luomus | Digital bone, claw and skin |
| 6291 | KN 6291 | *P. h. saimensis* | Full | 1971 | Finland: Lake Saimaa, Pyyvesi | U | Luomus | Digital bone, claw and skin |
| 6296 | KN 6296 | *P. h. saimensis* | Full | 1973 | Finland: Lake Saimaa, Kolovesi | U | Luomus | Digital bone, claw and skin |
| 6297 | KN 6297 | *P. h. saimensis* | Full | 1973 | Finland: Lake Saimaa, Heinävesi | F | Luomus | Digital bone |
| 6732 | KN 6732 | *P. h. saimensis* | Full | 1973 | Finland: Lake Saimaa, Heinävesi | F | Luomus | Digital bone and claw |
| 6368 | KN 6368 | *P. h. saimensis* | Full | 1975 | Finland: Lake Saimaa, Pihlajavesi | U | Luomus | Postcanine tooth |
| 6728 | KN 6728 | *P. h. saimensis* | Full | 1976 | Finland: Lake Saimaa, Pihlajavesi | F | Luomus | Digital bone and claw |
| 293 | 293 | *P. h. saimensis* | Full | 1981 | Finland: Lake Saimaa, Petraselkä | F | UEF | Muscle |
| 821 | 821 | *P. h. saimensis* | Full | 1985 | Finland: Lake Saimaa, Haukivesi | F | UEF | Muscle |
| 1394 | 1394 | *P. h. saimensis* | Full | 1991 | Finland: Lake Saimaa, Haukivesi | M | UEF | Muscle |
| 1687 | 1687 | *P. h. saimensis* | Full | 1996 | Finland: Lake Saimaa, Joutenvesi | F | UEF | Muscle |
| 2427 | 2427 | *P. h. saimensis* | Full | 2007 | Finland: Lake Saimaa, Pihlajavesi | M | UEF | Muscle |
| NN12-07 | NN12-07 | *P. h. botnica* | Full | 2008 | Finland: Baltic Sea, Bothnian Bay | M | LUKE | Muscle |
| NN8-07 | NN8-07 | *P. h. botnica* | Full | 2008 | Finland: Baltic Sea, Bothnian Bay | M | LUKE | Muscle |
| F = female, M = male, U = unknown sex, Luomus = The Finnish Museum of Natural History, UEF = University of Eastern Finland, LUKE = Natural Resources Institute Finland | | | | | | | | |
