## Supplementary Table S2 for "Museum specimens of a landlocked pinniped reveal recent loss of genetic diversity and unexpected population connections"

Supplementary Table S2. Primers used to amplify the mtDNA control region in ringed seals.

| **Name** | **Sequence 5’-3’** | **Primer used at** | **Reference** |
| --- | --- | --- | --- |
| H16305 | AAGGAAGAGGTAACAACCC | University of Oulu | Palo 2003 |
| R109 | CCAGGTGCGTGAATACTG | University of Oulu | This study |
| F108 | CCACCTTACTGTGCTATCA | University of Oulu | This study |
| R328 | TGATTTCCCGAGGCATGGTG | University of Oulu | This study |
| F322 | TCACTTAGTCCAAGAGCC | University of Oulu | This study |
| R471 | ATTTGAAGGGTTAGTAGGA | University of Oulu | This study |
| F425 | CTATACTGGAACTATATCT | University of Oulu | This study |
| R605 | GCGTCGAGACCTTTACG | University of Oulu | This study |
| F603 | AAGCGGTATCACTCAGCTATG | University of Oulu | This study |
| L224 | GTGTACGTAACGTAACTATGT | University of Oulu | Valtonen et al. 2012 |
| SL1F | GCACCCAAAGCTGACATTCT | Durham University | This study |
| SL1R | GCTTATATGCATGGGGCAAA | Durham University | This study |
| SL2F | ATCGTGCATTNAYGGTTTGC | Durham University | This study |
| SL2R | GTAYACGTTTCACAAGGGTTG | Durham University | This study |
| SL3F | ACCATGCCTCGGGAAATC | Durham University | This study |
| SL3R | GATCATGGNCTGNTTAGTCATT | Durham University | This study |
